# Single-sequence protein-RNA complex structure prediction by geometric attention-enabled pairing of biological language models

**DOI:** 10.1101/2024.07.27.605468

**Authors:** Rahmatullah Roche, Sumit Tarafder, Debswapna Bhattacharya

## Abstract

Ground-breaking progress has been made in structure prediction of biomolecular assemblies, including the recent breakthrough of AlphaFold 3. However, it remains challenging for AlphaFold 3 and other state-of-the-art deep learning-based methods to accurately predict protein-RNA complex structures, in part due to the limited availability of evolutionary and structural information related to protein-RNA interactions that are used as inputs to the existing approaches. Here, we introduce ProRNA3D-single, a new deep-learning framework for protein-RNA complex structure prediction with only single-sequence input. Using a novel geometric attention-enabled pairing of biological language models of protein and RNA, a previously unexplored avenue, ProRNA3D-single enables the prediction of interatomic protein-RNA interaction maps, which are then transformed into multi-scale geometric restraints for modeling 3D structures of protein-RNA complexes via geometry optimization. Benchmark tests show that ProRNA3D-single convincingly outperforms current state-of-the-art methods including AlphaFold 3, particularly when evolutionary information is limited; and exhibits remarkable robustness and performance resilience by attaining better accuracy with only single-sequence input than what most methods can achieve even with explicit evolutionary information. Freely available at https://github.com/Bhattacharya-Lab/ProRNA3D-single, ProRNA3D-single should be broadly useful for modeling 3D structures of protein-RNA complexes at scale, regardless of the availability of evolutionary information.

## Introduction

The interactions between protein and ribonucleic acid (RNA) underpin a wide range of cellular processes from gene regulation to transcription to protein synthesis (1-6), with enormous implications in understanding the molecular basis of disease (7-10). Therefore, knowledge of protein-RNA interactions in atomic detail is critically important. However, experimental determination of the atomic-level protein-RNA complex structures is not always feasible or practical (11,12). To address the challenges associated with experimental characterization of protein-RNA complexes, several computational methods for predicting the interaction between protein and RNA have been developed, including traditional methods based on template-based modeling (13,14) and protein-RNA docking (15-19). However, their predictive modeling performance remains limited by the availability of the homologous template information or the deficiencies of docking sampling and scoring.

Following the breakthrough of deep learning-based protein structure prediction methods AlphaFold2 (20) and RoseTTAFold (21), a handful of deep learning-based methods have been recently developed for structure prediction of biomolecular interactions including protein-RNA complex structure prediction. For example, Baker and co-workers extended the RoseTTAFold approach for the prediction of protein-nucleic acid complex structures by developing RoseTTAFold2NA (RF2NA) (22) and subsequently introduced a generalized biomolecular modeling and design framework by developing RoseTTAFold All-Atom (RF2AA) (23) that is capable of protein-RNA complex structure prediction. Very recently, Google DeepMind and Isomorphic Labs developed a new generation of the AlphaFold system, called AlphaFold 3 (AF3) (24), a single unified deep learning framework capable of predicting the joint structure of complexes of various biomolecular interactions including protein-RNA complexes. However, the prediction accuracies of even these state-of-the-art deep learning-based methods remain low for protein-RNA complex structure prediction. For example, the average iLDDT score (25) of AF3 is 39.4 for protein-RNA complex structure prediction task on the test set, whereas the average iLDDT of RF2NA is 19.0 (see Extended Data Table 1 of (24)). While AF3 clearly outperforms RF2NA, the overall accuracy of AF3 is far from accurate (<40%), indicating that there is a large room for improvement in predicting structures of protein-RNA complexes.

The limited successes in protein-RNA complex structure prediction are mainly due to: 1) lack of structural data in the Protein Data Bank (PDB) (26) beyond the most common families of RNA-binding proteins (RBPs) (27); and 2) even basic amino acid and nucleic acid joint information such as the evolutionary sequences of joint protein-RNA multiple sequence alignments (MSAs) are often shallow and lack sufficient coverage. Thus, it is worthwhile to develop single-sequence protein-RNA complex structure prediction methods.

Promisingly, protein language models (pLM) (28-32) pretrained on a large corpus of protein sequences using attention-based transformer neural network architectures (33) have shown unprecedented scalability and broad generalizability in a wide range of predictive modeling tasks— from protein structure prediction (32) to protein engineering (34,35). Alongside the progress in pLMs, significant advances have been made in pretrained RNA language models (36,37), showing promising performance in various downstream prediction tasks. These biological language models, pretrained with an unsupervised learning objective across a large corpus of sequence database, are highly effective for downstream structure-related tasks even if they are trained purely with sequence information or subsequently fine-tuned with small-scale structural data. Importantly, biological language models implicitly capture evolutionary patterns present in biological sequences by means of transformer’s attention mechanism, without the need to supply explicit evolutionary data in the form of MSAs. That is, an effective coupling of protein and RNA language models not only has the potential to compensate for the lack of structural data that only exist for the most common families of RBPs, but also reduces the dependence on the availability of explicit evolutionary information in the form of joint protein-RNA MSAs that are often scarce and noisy, paving the way to single-sequence prediction of protein-RNA complex structure. Given the progress, a natural question arises: can we develop an integrative modeling approach to combine a pair of biological language models – one pretrained using protein sequences and the other pretrained using RNA sequences – for single-sequence protein-RNA complex structure prediction with improved accuracy and robustness?

Here, we present ProRNA3D-single, a single-sequence protein-RNA complex structure prediction method by geometric attention-enabled pairing of biological language models. Instead of relying on explicit evolutionary information by means of joint protein-RNA MSAs, our single-sequence method directly leverages the biological language model embeddings for the input protein and RNA sequences together with single-sequence protein and RNA structure prediction using pretrained biological language models. By converting protein and RNA language model embeddings into a structure-aware graph representation, ProRNA3D-single employs symmetry-aware graph convolutions followed by a pairwise combination of the individual component embeddings to generate protein-RNA pair embedding. The pair embedding is then fed into a ResNet-Inception module to capture the inter-component interactions between the residue-nucleotide pairs, followed by a geometric attention module to account for the many-body effect in protein-RNA interactions while satisfying geometric consistency through an attention mechanism (33), which can predict the inter-component interactions, resulting in protein-RNA interaction maps. Finally, the predicted protein-RNA interactions are transformed into multi-scale geometric restraints to optimize the relative spatial position and orientation of the protein and RNA components, leading to protein-RNA complex structure prediction.

Our method ProRNA3D-single convincingly outperforms the state-of-the-art deep learning-based protein-RNA complex structure prediction methods, including RoseTTAFold2NA, RoseTTAFold All-Atom, and AlphaFold 3. Despite being a single-sequence method, our method attains improved accuracy compared to the MSA-dependent methods when MSA information is limited, and exhibits remarkable robustness and performance resilience by attaining better accuracy with only single-sequence input than what most state-of-the-art MSA-dependent methods can achieve even with explicit MSA information. We directly verify that the structural-level predictive modeling accuracy of our method is closely connected to the quality of the protein-RNA interaction maps predicted through geometric attention-enabled pairing of biological language models. The improved accuracy and robustness of ProRNA3D-single make it broadly useful for modeling 3D structures of protein-RNA complexes at scale, regardless of the availability of explicit evolutionary information. An open-source software implementation of ProRNA3D-single, licensed under the GNU General Public License v3, is freely available at https://github.com/Bhattacharya-Lab/ProRNA3D-single.

## Materials and methods

### Overview of ProRNA3D-single framework

**Fig 1** illustrates our ProRNA3D-single method for protein-RNA complex structure prediction. Different from the recent deep learning-based approaches for protein-RNA complex structure prediction that require explicit evolutionary information in the form of joint protein-RNA MSAs, our single-sequence method directly leverages the biological language model embeddings for the input protein and RNA sequences together with single-sequence protein and RNA structure prediction using pretrained biological language models. ProRNA3D-single first converts the protein and RNA language model embeddings into a structure-aware graph representation and then employ symmetry-aware graph convolutions followed by a pairwise combination of the individual component embeddings to generate protein-RNA pair embedding. The pair embedding is then fed into a ResNet-Inception module to capture the inter-component interactions between the residue-nucleotide pairs, followed by a geometric attention module to account for the many-body effect in protein-RNA interactions while satisfying geometric consistency through an attention mechanism (33), which can predict the inter-component interactions, resulting in protein-RNA interaction maps. Finally, the predicted protein-RNA interactions are transformed into multi-scale geometric restraints to optimize the relative spatial position and orientation of the protein and RNA components, leading to protein-RNA complex structure prediction.

**Fig 1.**
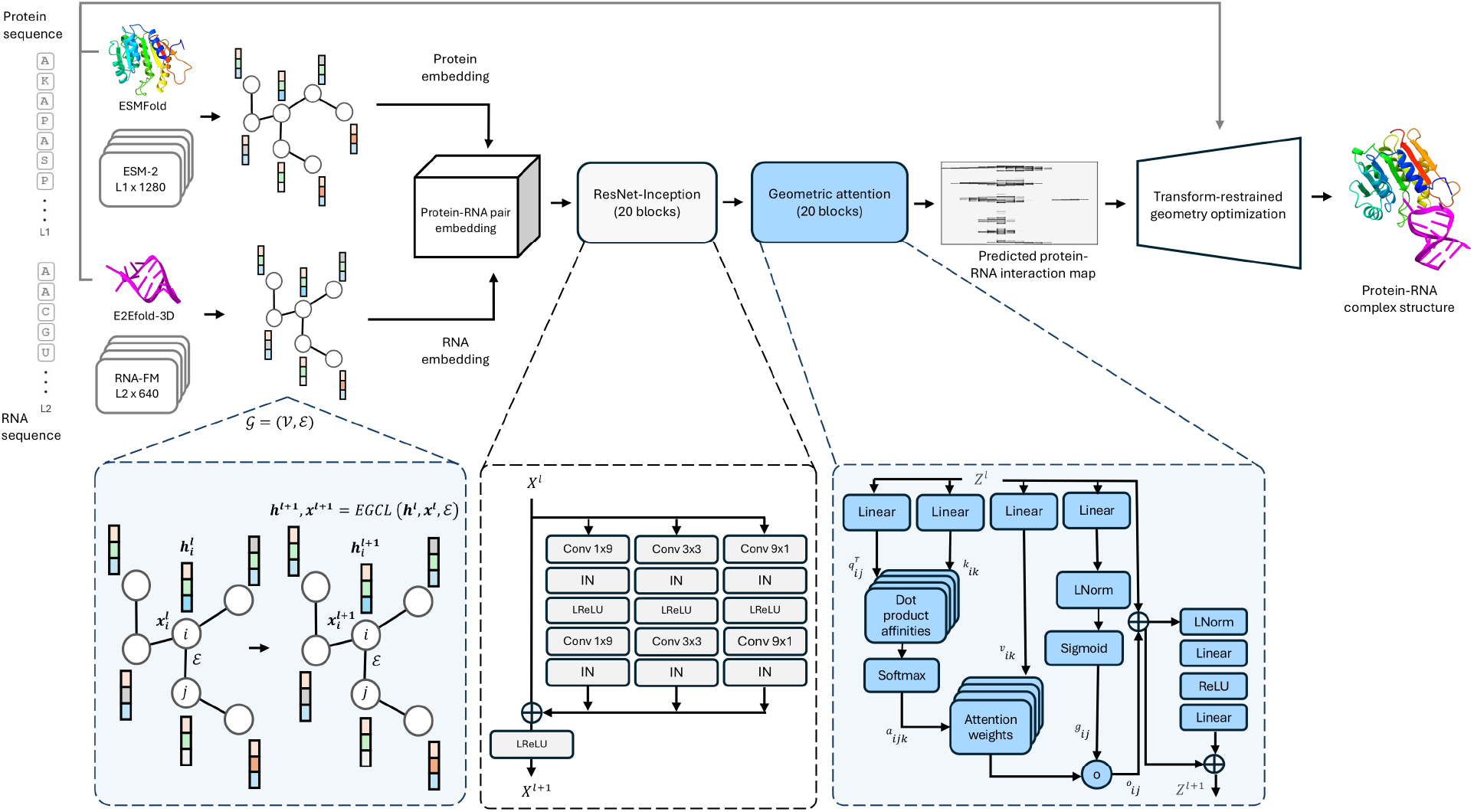
Illustration of ProRNA3D-single method for single-sequence protein-RNA complex structure prediction, consisting of three major modules. First, protein-RNA pair embedding generation using structure-aware graph representation of the biological language model embeddings for the input protein and RNA sequences, symmetry-aware graph convolutions, followed by a pairwise combination of the individual component embeddings. Second, protein-RNA interaction map prediction using a ResNet-Inception module followed by a geometric attention module. Third, protein-RNA complex structure prediction using transform-restrained geometry optimization.

### Protein-RNA pair embedding generation

Given the input protein and RNA sequences, we use ESMFold (32) to predict the structure of the input protein and E2EFold-3D (36) with ‘single sequence mode’ to predict the structure of the RNA. Both ESMFold and E2EFold-3D leverage pretrained biological language models for single-sequence structure prediction, without any explicit evolutionary information such as multiple sequence alignments (MSAs). In addition to single-sequence structure prediction, we generate language model embeddings with the input protein and RNA sequences using ESM-2 (32) and RNA-FM (37) models, respectively, with the protein embedding dimension of L1×1280 where L1 represents the length of the protein, and RNA embedding dimension of L2×640 where L2 represents the length of the RNA. The protein and RNA language model embeddings are then converted into two independent structure-aware graph representations *𝒢* = (*𝒱, ℰ*), where a node v ∈ *𝒱* contains the residue-level embeddings (for the protein graph) or the nucleotide-level embeddings (for the RNA graph) as the node features, and an edge *e* ∈ *ℰ* represents an interacting residue pair (for the protein graph) or a nucleotide pair (for the RNA graph). A residue pair is considered to be interacting if the Euclidean distance between the corresponding C_α_ atoms is within 14Å; and a nucleotide pair is considered to be interacting if the Euclidean distance between the corresponding C’_4_ atoms is within 20Å. The distance thresholds are chosen following the previous studies (38-40). For the edge features in both protein and RNA graph representations, we utilize the ratio of *log* |*i-j*|, and ||*i-j*||^2^, where |*i-j*| *i*s the absolute sequence difference between two residues (for protein) or two nucleotides (for RNA), and ||*i-j*||^2^ is the Euclidean distance between the residues (or nucleotides). In addition, we use coordinate information, extracted directly from the Cartesian coordinates of the C_α_ atoms of the predicted protein structures and the C’_4_ atoms of the predicted RNA structures.

The node and edge features together with the coordinate information are then fed into separate protein- and RNA-specific E(3)-equivariant graph neural networks (EGNNs) (41) to employ symmetry-aware graph convolutions. Both the protein- and RNA-specific EGNNs consist of a series of equivariant graph convolution layers (EGCL) to perform a series of transformations to the inputs. The EGCL updates the node and coordinate embeddings based on the node, coordinate, and edge embeddings from the previous layer, eventually resulting in enriched protein and RNA embeddings 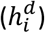 by applying a linear transformation to the last layer of the protein- and RNA-specific EGNNs. Formally, the EGCL operations are defined as follows:

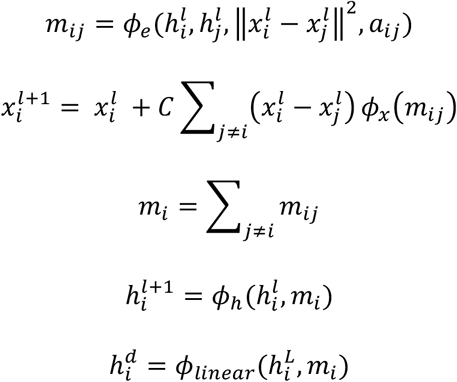

where *ϕ*_*e*_, *ϕ*_*x*_ and *ϕ*_*h*_ are non-linear transformations for edge (*m*_*ij*_), coordinate 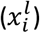 and node 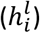 embeddings at node *i* in layer *l*; *m*_*i*_ is the aggregated message at node *i*, and *ϕ*_*linear*._ represents the linear transformation on the final layer node embeddings 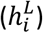 to obtain the enriched embeddings 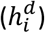. Both the protein- and RNA-specific EGNNs are built on 4 layers of EGCL with a hidden dimension of 128. The number of layers and hidden dimensions are determined using an independent validation set (see **Supplementary Table 1**). Finally, a pairwise combination is performed between the enriched protein embeddings having a dimension of (L1×D) where L1 represents the length of the protein, and the enriched RNA embeddings having a dimension of (L2xD) where L2 represents the length of the RNA, resulting in protein-RNA pair embedding having a dimension of (L1×L2×2D).

### Protein-RNA interaction map prediction

The protein-RNA pair embedding is passed as input to a two-stage deep learning architecture consisting of a ResNet-Inception module followed by a geometric attention module to predict inter-protein-RNA interaction maps. The ResNet-Inception module aims to capture the inter-component interactions between the residue-nucleotide pairs through a neural architecture comprising of a stack of residual neural networks (ResNet) having three parallel branches of convolutional layers (42,43). The three parallel branches each consists of two convolutional layers with kernel size of (1×9), (3×3), and (9,1), respectively. In each branch, an instance normalization followed by a Leaky ReLU (44) is applied to the output of the first convolutional layer, and the output of the activation layer is passed to the second convolutional layer. A shortcut connection is applied from the input to the output of all the three branches, and the final output followed by a Leaky ReLU activation is passed to the next ResNet-Inception block as input. Formally, the protein-RNA pair embedding is updated through the ResNet-Inception block as follows:

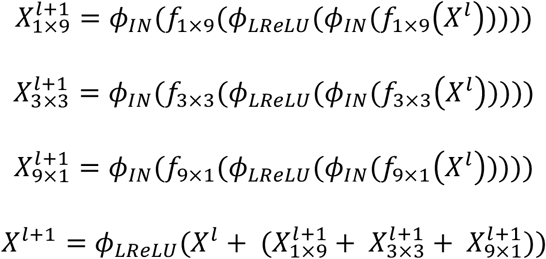

where, *X*^*l*^ and *X*^*l+*1^are the pair representation at layer *l* and *l+1*, respectively, *ϕ*_*IN*_ is the instance normalization, *ϕ*_*LReLU*_is the Leaky ReLU activation operation. The 2D convolutions with kernel size of (1×9), (3×3), and (9×1) are represented by *f*_1×9_, *f*_3×3_, and *f*_9×1_, respectively. The network architecture of the ResNet-Inception module consists of 20 consecutive blocks, determined using an independent validation set (See **Supplementary Table 1**).

To account for the many-body effect in protein-RNA interactions while satisfying geometric consistency, a geometric attention module is subsequently employed that performs triangle-aware self-attention mechanism similar to a recent study (43). Formally, the triangle-aware self-attention mechanism is defined as follows:

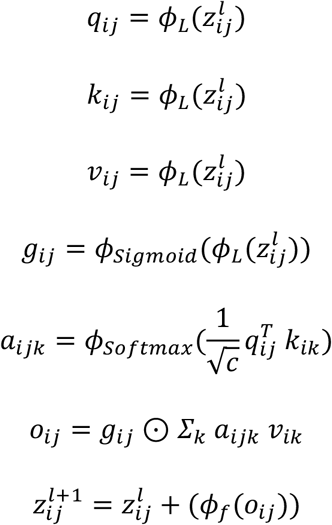

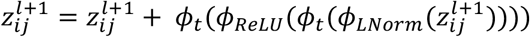

where, *ϕ*_*L*_ is the linear operation that transforms the triangle-aware pair representation 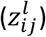 at layer *l* into query (*q*_*ij*_), key (*k*_*ij*_), and value (*v*_*ij*_) representations for a pair (*i, j*). An additional gate (*g*_*ij*_) representation is obtained from a linear transformation followed by a non-linear sigmoidal operation. The multi-head attention weight (*a*_*ijk*_) between the three points (*i, j, k*) in the triangle is utilized to obtain the updated representation (*o*_*ij*_), which is transformed through a final linear operation (*ϕ* _*f*_). Finally, a non-linear transition transformation involving a layer normalization (*ϕ*_*LNorm*_), ReLU Activision (*ϕ*_*ReLU*_), and two linear operations (*ϕ*_*t*_) is applied along with short-cut connection to obtain the updated triangle-aware pair representation 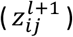. The network architecture of the geometric attention module consists of 20 triange-aware self-attention blocks, determined using an independent validation set (See **Supplementary Table 1**).

The output from the geometric attention module undergoes a layer normalization, followed by a 1×1 convolutional layer for predicting the pairwise distance between the C_α_ atom of a protein residue and the C4’ atom of an RNA nucleotide, resulting in a protein-RNA interaction map. The predicted distances are distributed across 37 bins, ranging from 2.5 Å to 20 Å with a step size of 0.5 Å, resulting in 36 bins. The final (37th) bin is considered for the distance exceeding 20 Å.

### Training details

Our method is implemented using Pytorch 1.12.0 (45). During model training, we use cross entropy loss function and ADAM (46) optimizer with a learning rate of 1e-4. Additionally, cosine annealing from SGDR (47) and a weight decay of 1e-16 are used. The model training runs for at most 100 epochs on a single NVIDIA-A100 GPU. In addition to the full-fledged version of our method, we separately train multiple baseline models by discarding equivariant updates (thus making the model invariant), by replacing the protein- and RNA-specific EGNNs with convolutional blocks for protein-RNA pair embedding generation; and by discarding geometric attention module employing triangle-aware self-attention.

### Transform-restrained geometry optimization

The predicted inter-protein-RNA interaction maps are transformed into multi-scale geometric restrains to optimize the relative spatial position and orientation of the protein and RNA components using PyRosetta-based relaxation algorithm (48). First, we transform the predicted distance maps into hybrid interaction maps with 10-level thresholding, similar to previous studies (49,50). Specifically, we obtain a hybrid interaction map with upper limits set at 2.5 Å, 4 Å, 6 Å, 8 Å, 10 Å, 12 Å, 14 Å, 16 Å, 18 Å, and 20 Å by summing the predicted likelihood values up to the respective distance thresholds. Subsequently, we generate ‘BOUNDED’ ‘AtomPair’ constraints in PyRosetta for the interactions having likelihood values ≥ 0.5 and utilize scaler-weighted function where the predicted likelihood values are incorporated as the scaler weights the lower bounds of the restraints are set to 2 Å, and the upper bounds are derived from the hybrid interaction maps. Finally, we feed the restraints to the Rosetta FastRelax algorithm (51) implemented in PyRosetta (48), which employs gradient-based optimization to generate 3D complex protein-RNA structural models. The initial pose originates from the previously described single-sequence protein and RNA monomers generated by language model-based ESMFold and E2Efold-3D, respectively. We set the following parameters for initialization ‘-hb_cen_soft -constant_seed -relax:default_repeats 5 - default_max_cycles 200 -out:level 100’, and utilize ‘ref2015’ as the scoring function and a maximum FastRelax iteration of 500.

### Datasets, competing methods, and performance evaluation

To train our new method, ProRNA3D-single, we collect X-ray crystal structures of protein-RNA complexes having resolution up to 3.5 Å that are deposited to the PDB on and before October 2022. Furthermore, the redundancy among the collected structures is removed following previous studies (52,53). We also discard structures with a) proteins chain length>1500, b) RNA chain length>500, and c) RNA chain length<10. Finally, we obtain a total of 798 non-redundant interacting protein-RNA complex structures, from which we consider 750 targets for training and 48 targets for validation set.

Using protein-RNA complexes published more recently than any training- and validation-set targets, we evaluate the predictive modeling accuracy of our method. Our test set consists of targets released on and after November 2022 till November 2023. Following a recent study (54), we further filter the test targets ensuring that test protein chains do not share over 50% sequence identity with the train and validation set using CD-HIT (55) to reduce their overlap, resulting in a total of 39 protein-RNA complexes for the test set, denoted by Test_39 in our performance benchmarking.

We compare our method with three state-of-the-art methods for predicting biomolecular interactions in general and protein-RNA complex structures in particular, including RosettaFold2NA (RF2NA) (22), RosettaFold All Atom (RF2AA) (23), and AlphaFold 3 (AF3) (24). RF2NA predicts multimeric protein, RNA, DNA and their complexes by utilizing MSA information for protein, RNA as well as the paired protein/RNA using a 3-track architecture introduced in RoseTTAFold (21) with 1D-track utilizing multiple sequence alignments (MSAs), 2D-track utilizing pairwise distances, and 3D-track utilizing structural information. In RF2NA, the 1D-track is extended from RoseTTAFold by including the additional input tokens for nucleotides, 2D-track is extended by including the interaction between nucleotides and amino acids, and the 3D-track is extended by including the coordinate frames and 10 torsion angles for backbone and side chains for nucleic acids. RF2AA is a generalized biomolecular modeling and design system that includes the prediction of protein-RNA complex structures. RF2AA system represents arbitrary molecules as atom-bond graphs that initially are disconnected gas of amino acid residues, nucleic acid bases, and freely moving atoms, which are successively transformed through multiple blocks of the 3-track network architecture into physically plausible assembled structures. AlphaFold 3 is a unified deep learning framework for predicting biomolecular interactions that is substantially updated from AlphaFold2 (20) by introducing a lightweight Pairformer module for MSA processing and a diffusion-based architecture for directly predicting atomic coordinates starting from the single chain to complex structures including protein, nucleic acids, and small ligands. While RF2AA and RF2NA are publicly available methods, the code and model weights of the AlphaFold 3 are not yet publicly available, except for a webserver (alphafoldserver.com) with limited job submission and restrictions on its usage. Therefore, we obtain the predictions by AlphaFold 3 (AF3) by directly submitting the jobs in the alphafoldserver.com webserver and for the comparison, we consider the best model (model 0) out of 5 predictions based on its default self-estimation scores. The RF2NA and RF2AA predictions are obtained by running them locally, after their inference code and model weights are downloaded in October 2023, and in April 2024, respectively. It is worth noting that all three competing state-of-the-art methods utilizes MSA and template information for prediction, whereas our new method, ProRNA3D-single is a single-sequence method. Therefore, for a fair comparison with our template- and MSA-free method ProRNA3D-single, we additionally run a customized version of RF2NA without MSA and template information, naming it RF2NA-single.

To evaluate the accuracy of the protein-RNA 3D complex structures, we use fnat score, a widely used CAPRI metric (56) for evaluating the interface of predicted biomolecular interactions. The metric fnat denotes the fraction of native contacts that are present in the predicted structure. In the case of protein-RNA interactions, a residue-nucleotide pair is considered to be in contact if their heavy atoms are within 5 Å. The fnat scores lie in range of [0-1], and the higher score represents better accuracy of the predicted complex structural model. We use two additional metrics named iRMS and lRMS for the evaluation of the predicted protein-RNA complex structures. The score for iRMS (interface root mean square) is relatively relaxed metric, where the interface is defined by an atomic contact cut-off of 10 Å, and the root mean square deviations of the backbone atoms are calculated after optimally superposing them with the native. The third metric lRMS (ligand root mean square) is measured by calculating the root mean square deviations of the shorter chain backbone atoms after optimally superposing the longer chain with the native. For both iRMS and lRMS, lower scores represent better accuracy of the predicted structural models. We use DockQ program (57) (downloaded in December 2023) for measuring the fnat, iRMS, and lRMS metrics.

## Results

### Test Set Performance

**Fig 2** shows the predictive modeling accuracy of our new method ProRNA3D-single compared to the state-of-the-art deep learning-based methods AF3, RF2AA and RF2NA on the test set, Test_39. ProRNA3D-single attains the best fnat scores compared to the state-of-the-art methods, attaining 41.03% “successful prediction” having fnat>0.2, which is 25.64%, 25.64%, and 17.95% higher than RF2NA, RF2AA, and AF3, respectively (**Fig 2 B**), where a prediction is considered “successful” if the predicted structural model has fnat>0.2 following CAPRI criteria and previous studies (56,58). ProRNA3D-single also attains better performance compared to the competing methods in terms of iRMS **(Fig 2 C)** and lRMS **(Fig 2 D)**. Of note, ProRNA3D-single is the only single-sequence method, whereas all competing methods take advantage of explicit evolutionary information in the form of MSAs as well as homologous templates whenever available. That is, ProRNA3D-single attains state-of-the-art performance despite being a single-sequence method. When compared head-to-head against the customized version of RF2NA without MSA and template information, RF2NA-single, which fails to predict any model with fnat>0.2, our method ProRNA3D-single outperforms across all metrics by a large margin.

**Fig 2.**
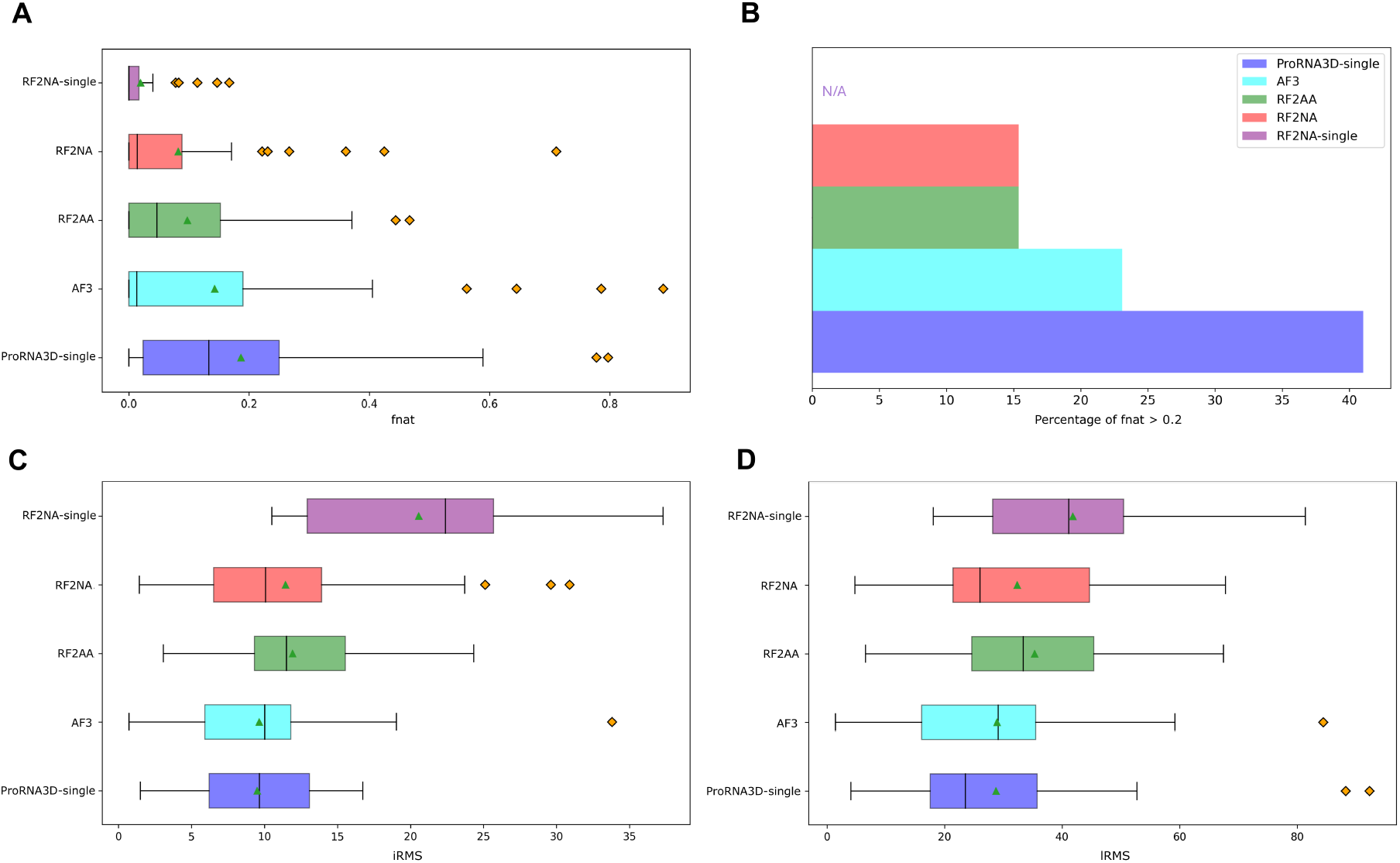
Test set performance of ProRNA3D-single and the competing methods. (**A**) The distributions of the fnat scores. (**B**) Percentage of “successful predictions” having fnat>0.2. (**C**) The distributions of the iRMS scores. (**D**) The distributions of the lRMS scores.

A representative example from the test dataset shown in **Fig 3** illustrates the advantage of ProRNA3D-single over the competing methods. This example, representing TRANSFERASE/RNA of Enterovirus A71 (PDB ID 7W9S, protein chain A, RNA chain C), falls in a distinct group of nucleic acid polymerases encoded by RNA viruses, which are crucial for viral genome replication and transcription, making them prime candidate for the development of antiviral drugs (59). Our method yields an accurate prediction attaining fnat score of 0.778 that is much better than AF3, RF2AA, and RF2NA. It is interesting to note that for this target, all competing methods except RF2NA fail to predict structures with fnat>0.2, with AF3 delivering the lowest performance. The performance advantage of ProRNA3D-single is striking with 0.778, 0.75, and 0.556 in fnat score points gain for ProRNA3D compared to AF3, RF2AA, and RF2NA, respectively.

**Fig 3.**
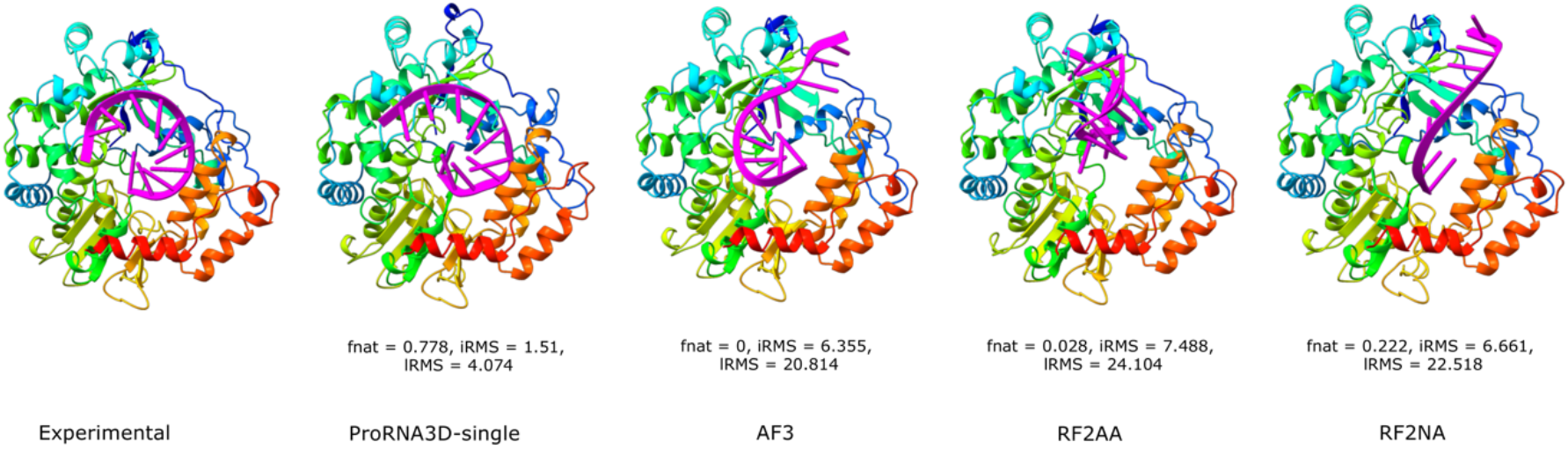
A representative example of protein-RNA complex structure prediction using ProRNA3D-single and the competing methods for PDB ID 7w9s, protein chain A, RNA chain C, representing TRANSFERASE/RNA of Enterovirus A71. Protein molecules are rainbow colored from blue to red from the N-to C-termini and RNA molecules are colored in magenta.

### Effect of evolutionary sequence information

While our new method, ProRNA3D-single, is a single-sequence method harnessing the power of biological language models, without utilizing any explicit evolutionary sequence information in the form of MSAs (or homologous templates), the competing methods rely on the availability of explicit MSA information (and homologous templates whenever available). To investigate how the availability of explicit evolutionary sequences in the form of protein/RNA paired MSAs affects the prediction accuracy of the competing methods compared to the MSA-agnostic ProRNA3D-single, we split the Test_39 set into two subgroups based on the paired MSAs generated using the MSA search engine of RF2NA : one where the MSA search is unsuccessful and there are no evolutionary sequences in paired MSA except for the query sequence (paired MSA depth = 1), and the other when there exists evolutionary sequences in paired MSA (paired MSA depth > 1). As shown in **Fig 4**, 28 out of 39 targets in the test set, the MSA search is unsuccessful where the protein/RNA paired MSA depth is 1. That is, for the majority of targets, there is no protein/RNA paired MSA, highlighting the scarcity of explicit evolutionary information and the urgent need to develop single-sequence methods for protein-RNA complex structure prediction. For the subset of targets where the paired MSA depth = 1, our single-sequence method ProRNA3D-single attains remarkably better accuracy than AF3, RF2NA, and RF2AA (**Fig 4 inset A, Supplementary Table 2**). For the subset of targets when there exist evolutionary sequences (paired MSA depth > 1), ProRNA3D-single still attains state-of-the-art performance, second only to AF3 in terms of fnat score, while outperforming RF2AA and RF2NA (**Fig 4 inset B, Supplementary Table 2**), thereby exhibiting performance resilience. It is interesting to note that AF3 and RF2NA attain higher overall accuracy for the subset of targets having explicit evolutionary information (paired MSA depth > 1) compared to the subset where there are no evolutionary sequences in paired MSA except for the query sequence (paired MSA depth = 1), demonstrating that the availability of (or lack thereof) evolutionary sequences affects the predictive modeling accuracy of the MSA-dependent methods. In contrast, the prediction accuracy of our single-sequence method ProRNA3D-single, remains unaffected by explicit evolutionary information, demonstrating its robustness.

**Fig 4.**
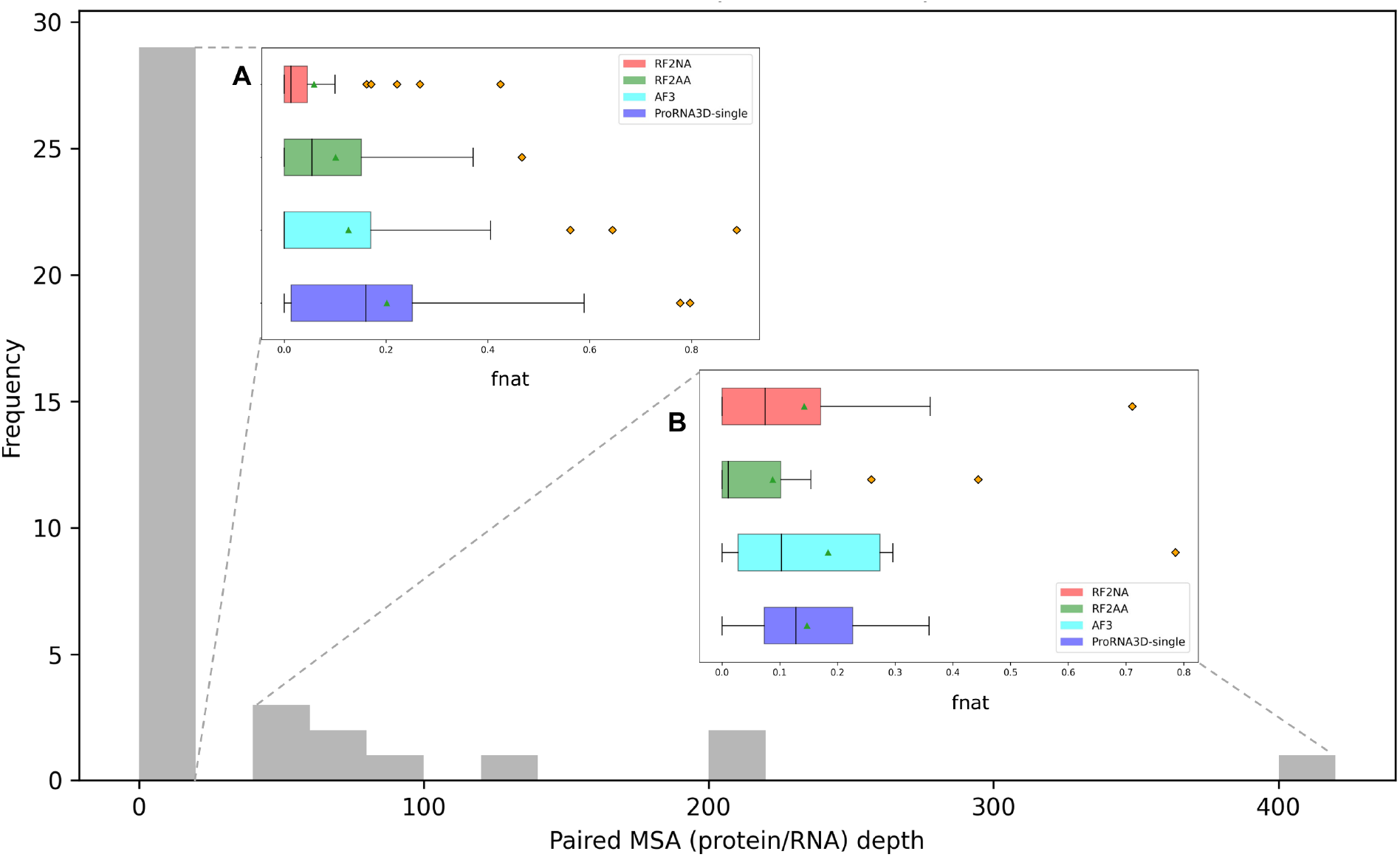
The distribution of evolutionary sequences in the Test_39 set is represented as the histogram of protein/RNA paired MSAs generated using the evolutionary sequence search engine of RF2NA. (**Inset A**) Performance of ProRNA3D-single and the competing methods when there is no sequence in paired MSA except for the query sequence (paired MSA depth = 1). (**Inset B**) Performance of ProRNA3D-single and the competing methods when there exist evolutionary sequences in paired MSA (paired MSA depth > 1).

Two representative examples shown in **Fig 5** may help further shed light on the effect of evolutionary sequences. The first example, HYDROLEASE/RNA for Macaca mulatta (target ID 7UU3, protein chain B and RNA chain D), is from the subset of targets where there are no evolutionary sequences in paired MSA except for the query sequence (paired MSA depth = 1). ProRNA3D-single attains much better accuracy compared to the competing methods with 0.457, 0.457, and 0.086 in fnat score points gain for ProRNA3D-single compared AF3, RF2NA and RF2AA, respectively. The improved accuracy of ProRNA3D-single is visually noticeable, whereas AF3 and RF2NA fail to predict accurate protein-RNA interface and RF2AA suffers from unrealistic RNA conformation. The second example of RNA Binding Protein/RNA for Homo Sapiens (target ID 8ID2, protein chain A, RNA chain C) is from the subset having explicit evolutionary information (43 sequences in paired MSA). ProRNA3D-single still outperforms AF3, RF2NA and RF2AA by 0.077, 0.282, and 0.205 fnat score points, respectively. In both cases, our method consistently yields “successful prediction” having fnat>0.2, whereas only RF2AA in the first example and AF3 in the second example attain fnat>0.2 among the three competing methods.

**Fig 5.**
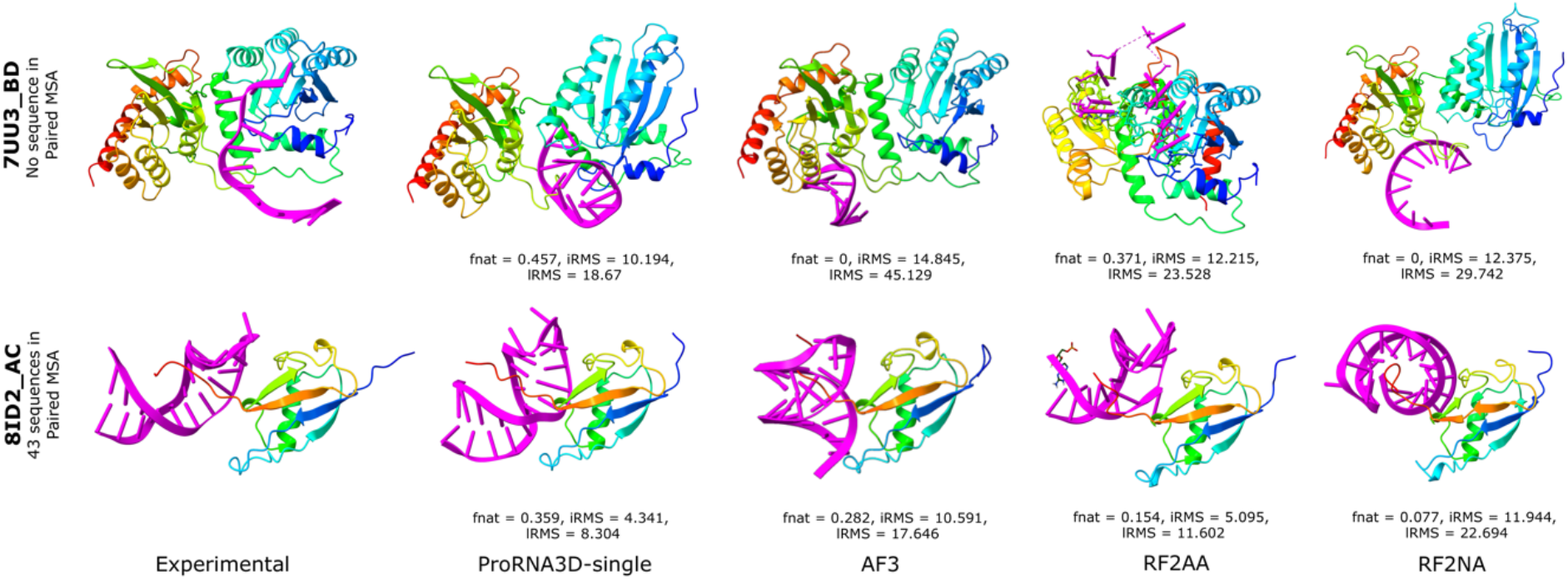
Protein-RNA complex structure prediction by ProRNA3D-single and the competing methods for two representative examples: HYDROLEASE/RNA for Macaca mulatta (PDB ID 7UU3, protein chain B and RNA chain D), and RNA Binding Protein/RNA for Homo Sapiens (PDB ID 8ID2, protein chain A, RNA chain C).

### Contribution of the interatomic protein-RNA interaction map prediction

A core component of our method is the prediction of interatomic (C_α_ - C’_4_) protein-RNA interaction maps, which are subsequently transformed into geometric restraints to optimize the relative spatial position and orientation of the protein and RNA components, leading to protein-RNA complex structure prediction. Therefore, a natural question to ask is, does the quality of the predicted interaction maps translate into a good (or bad) complex structure prediction? To examine the contribution of interatomic protein-RNA interaction map prediction, we evaluate the precision of contact maps extracted from the hybrid interaction maps with 10-level thresholding in the Test_39 set. **Fig 6 A** represents the average precision of the predicted contacts at the terminal distance threshold of 20 Å. Considering the top1, top10, and top predicted contacts having likelihood values ≥ 0.5 at 20 Å threshold, our interatomic (C_α_ - C’_4_) protein-RNA interaction predictions have the average precision of 43.59%, 35.385%, and 35.374% respectively (a full set of the average precision scores can be found in **Supplementary Table 3**). More importantly, the precision of the interaction prediction maintains a high correlation (Pearson r=0.75) with the fnat scores of the resulting complex structures of the corresponding targets (**Fig 6 B)**, verifying that the structural-level predictive modeling accuracy of our method is closely connected to the quality of the protein-RNA interaction maps.

**Fig 6.**
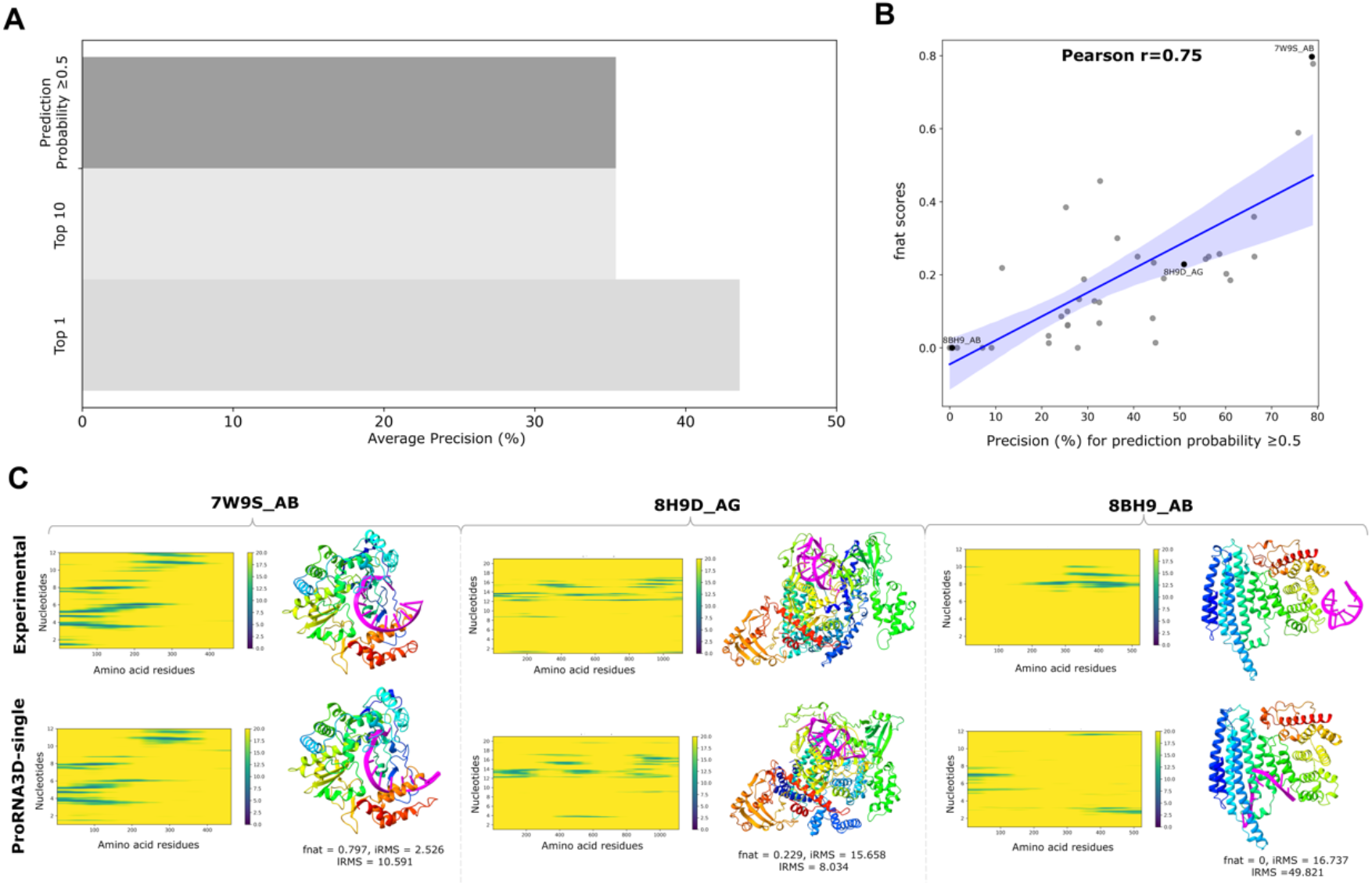
(**A**) The interatomic (C_α_ - C’_4_) protein-RNA interaction prediction performance at 20 Å thresholds. Average precision for top 1, 10, and interactions with prediction probability ≥ 0.5 are reported. (**B**) Scatterplot of precision (%) for prediction probability ≥ 0.5 at 20 Å thresholds against the fnat scores in the Test_39 set. (**C**) Interatomic (C_α_ - C’_4_) protein-RNA interaction maps predicted by our method compared to the native distance maps along with the corresponding predicted and native structures for three representative examples: TRANSFERASE/RNA of Enterovirus A71 (PDB ID 7W9S, protein chain A, RNA chain B), DNA binding protein of Lachnospiraceae bacterium (PDB ID 8H9D, protein chain A, RNA chain G), and RNA binding protein of Candida tropicalis MYA-3404 (PDB ID 8BH9, protein chain A, RNA chain B).

**Fig 6 C** shows three representative examples of our predicted interactions and the resulting 3D structural models, compared to the corresponding ground truth (experimental) distance maps and structures. For the TRANSFERASE/RNA of Enterovirus A71 (target ID 7W9S, protein chain A, RNA chain B), the interactions predicted by our method exhibit a clear similarity with the corresponding ground truth distance map, and the resulting 3D structural model demonstrates high accuracy, achieving an fnat score of 0.797. For the DNA binding protein of Lachnospiraceae bacterium (target ID 8H9D, protein chain A, RNA chain G), the predicted interactions are somewhat similar to the ground truth distance map, and the resulting structural model is deemed as an acceptable prediction with an fnat score of 0.229. Lastly, for RNA binding protein of Candida tropicalis MYA-3404 (target ID 8BH9, protein chain A, RNA chain B), our method yields inaccurate interaction prediction, showing significant dissimilarity to the corresponding ground truth distance map. Consequently, the resulting structural model is inaccurate. Overall, our analysis collectively demonstrates that the quality of predicted interactions is a key driver of the structural-level predictive modeling accuracy of our method, underscoring the importance of accurate inter-protein-RNA interaction prediction for improved protein-RNA complex structure prediction.

### Ablation study

Recognizing the importance of the interatomic (C_α_ - C’_4_) protein-RNA interaction prediction of our method, we investigate the relative contribution of different components of our interaction prediction networks. For that purpose, we conduct an ablation study to separately train multiple baseline models on the same train set by discarding equivariant updates (w/o equivariance) making the model invariant, by replacing the protein- and RNA-specific EGNNs with convolutional blocks for protein-RNA pair embedding generation (w/o graph embedding); and by discarding geometric attention module employing triangle-aware self-attention (w/o geometric attention) and compare their performance head-to-head with the full-fledged version of ProRNA3D-single on the Test_39 set.

**Fig 7** shows the performance comparison of ProRNA3D-single full-fledged model and the ablated baseline models. The average fnat scores of model w/o geometric attention, model w/o equivariance, and w/o graph embedding are 0.125, 0.116, and 0.123, noticeably lower than that of ProRNA3D-single default (0.186). Furthermore, the ablated baseline models exhibit reduced consistency having 17.95%, 28.21%, and 17.95% lower “successful prediction” having fnat>0.2 than the full-fledged ProRNA3D. Overall, the results demonstrate the performance benefits of employing symmetry-aware graph convolutions and geometric attention module, which are the key modules of the neural architecture of ProRNA3D-single.

**Fig 7.**
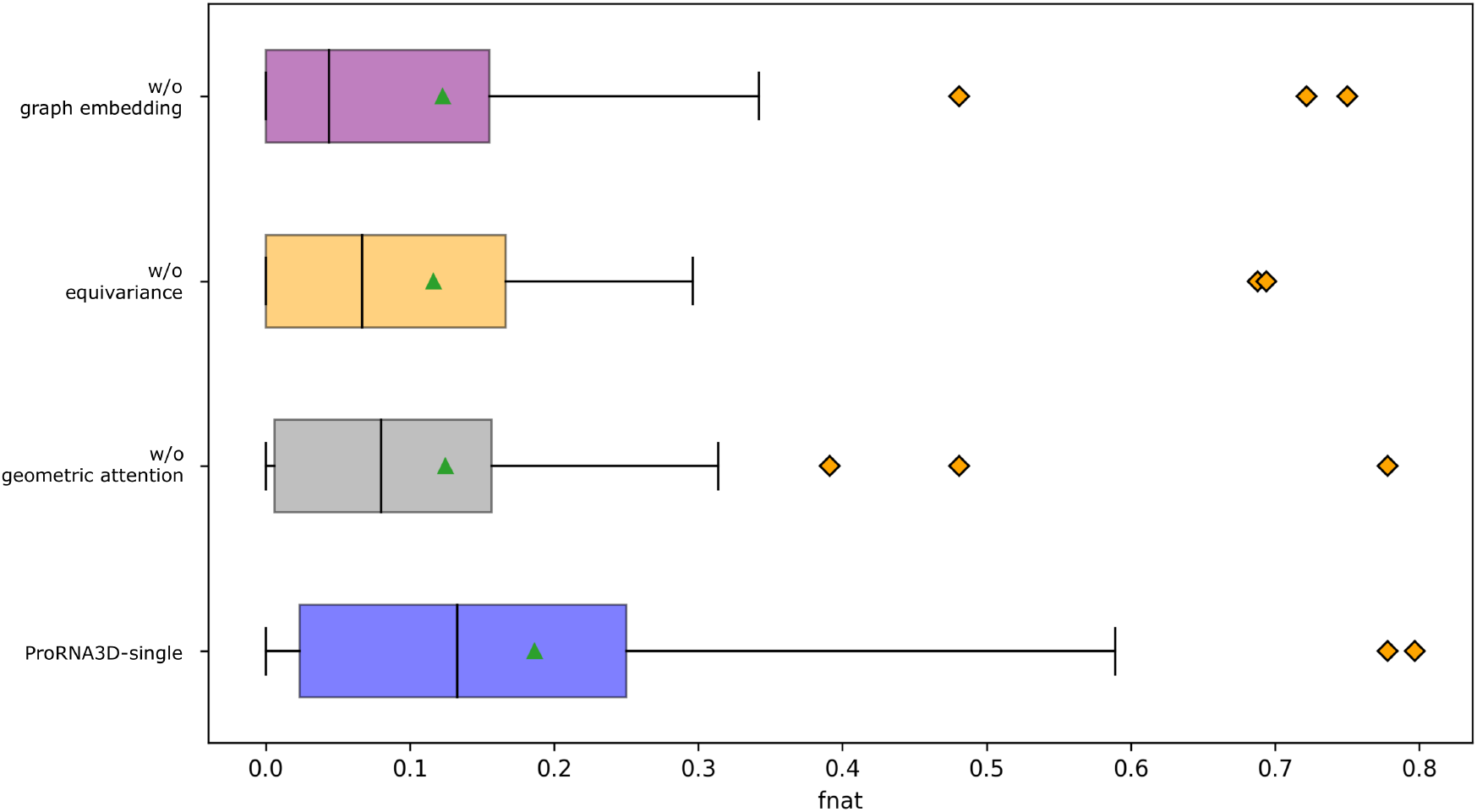
Ablation study showing the fnat distributions of ProRNA3D-single and the ablated baseline models on the Test_39 set.

## Discussion

This work introduces ProRNA3D-single, a new single-sequence method for predicting protein-RNA complex structures powered by biological language models. We demonstrate that ProRNA3D-single consistently outperforms the state-of-the-art deep learning-based protein-RNA complex structure prediction methods, including RoseTTAFold2NA, RoseTTAFold All-Atom, and AlphaFold 3. A major contribution of our work is the development of a generalized neural architecture for pairing biological language model embeddings, a previously unexplored avenue in the context of predicting biomolecular interactions. Experimental results show that our method attains substantially improved accuracy compared to the MSA-dependent methods when MSA information is limited, and exhibits remarkable robustness and performance resilience by attaining better accuracy in single-sequence protein-RNA complex structure prediction than what most state-of-the-art MSA-dependent methods can achieve even with explicit MSA information. Bypassing the dependence on the availability of explicit evolutionary information, which is not always abundant, our method paves the way to scalable and generalizable modeling of protein-RNA complex structures. Our ablation study reveals that the interplay of various modules in our neural network architecture cooperatively contributes to the improved accuracy, and the structural-level predictive modeling accuracy is closely connected to the quality of the protein-RNA interaction maps predicted through geometric attention-enabled pairing of biological language models.

Despite the improved performance of our method, there is still substantial room for improvement in protein-RNA complex structure prediction. A promising direction for future work is to investigate the possibility of incorporating biophysical “background knowledge” and inclusion of high-throughput experimental data in training to compensate for the lack of sufficient high-resolution structural data for protein-RNA complexes. Furthermore, we will study whether 3D modeling accuracy can be further improved through an integrative modeling approach that combines multimodal experimental data with deep learning-based predicted geometric restraints. Finally, while we find that ProRNA3D-single exhibits excellent predictive accuracy and remarkable robustness by geometric attention-enabled pairing of biological language models, an open challenge that remains is how to make the biological language models “partner-aware” specifically for biomolecular interaction prediction tasks. To this end, the possibility of fine-tuning biological language models using partner-specific knowledge may be explored. We expect our proposed method can be easily extended to predict the complex structures of other biomolecular interactions, including protein-DNA and protein-ligand as well as molecular assemblies containing proteins, small molecules, metals, and chemical modifications with improved accuracy and robustness.

## Availability

An open-source software implementation of ProRNA3D-single, licensed under the GNU General Public License v3, is freely available at https://github.com/Bhattacharya-Lab/ProRNA3D-single.

## Acknowledgements

This work was made possible in part by a grant of high-performance computing resources and technical support from the National AI Research Resource Pilot award (NAIRR240093 to D.B.).

## Funding

This work was partially supported by the National Institute of General Medical Sciences (R35GM138146 to D.B.) and the National Science Foundation (DBI2208679 to D.B.).

## Notes

### Competing Interest Statement

The authors have declared no competing interest.

