## Supplementary Information for "Single-sequence protein-RNA complex structure prediction by geometric attention-enabled pairing of biological language models"

**Supplementary Table 1.** Hyperparameter selection on the validation set. The average precision of top 1 and top 10 contacts at 20Å are reported. Values in bold represent the hyperparameters selected in ProRNA3D-single.

| **Hyperparameters** | | **top 1** | **top 10** |
|---|---|---|---|
| #of equivariant graph convolution layers | 2 | 29.167 | 28.958 |
| | **4** | 39.583 | 37.917 |
| | 6 | 25 | 26.875 |
| | 8 | 27.083 | 25.833 |
| Hidden dimension for equivariant graph convolution | 64 | 27.083 | 28.125 |
| | **128** | 39.583 | 37.917 |
| | 256 | 18.75 | 27.292 |
| # of ResNet +Triangle-aware attention blocks | 10 | 22.917 | 17.292 |
| | 15 | 33.333 | 31.042 |
| | **20** | 39.583 | 37.917 |
| | 25 | 18.75 | 26.667 |

**Supplementary Table 2.** Performance comparison of our method with AF3, RF2NA, and RF2AA in terms of average fnat, iRMS, and lRMS on the two subgroups of the Test_39 set based on the paired protein-RNA-MSA depth. Values in bold represent the best performance.

| **Benchmarking groups** | **Methods** | **fnat** | **iRMS** | **lRMS** |
|---|---|---|---|---|
| Paired protein-RNA-MSA depth = 1 | ProRNA3D-single | **0.202** | **8.581** | **23.562** |
| | AF3 | 0.126 | 9.411 | 28.616 |
| | RF2NA | 0.058 | 9.748 | 28.131 |
| | RF2AA | 0.101 | 10.801 | 32.412 |
| Paired protein-RNA-MSA depth > 1 | ProRNA3D-single | 0.147 | 11.827 | 41.882 |
| | AF3 | **0.184** | **10.243** | **29.737** |
| | RF2NA | 0.142 | 15.716 | 43.128 |
| | RF2AA | 0.088 | 14.811 | 42.695 |

**Supplementary Table 3.** The interatomic ($C_\alpha$ - $C'_4$) protein-RNA interaction prediction performance at different distance thresholds. Average precision for top 1, 10, and top predicted interactions having likelihood values ≥ 0.5 are reported.

| Thresholds | 2.5 Å | 4 Å | 6 Å | 8 Å | 10 Å | 12 Å | 14 Å | 16 Å | 18 Å | 20 Å |
|---|---|---|---|---|---|---|---|---|---|---|
| **Top 1** | 0.000 | 2.564 | 2.564 | 5.128 | 7.692 | 23.077 | 28.205 | 33.333 | 46.154 | 43.59 |
| **Top 10** | 0.000 | 0.513 | 3.077 | 7.179 | 12.308 | 20.769 | 27.179 | 33.077 | 36.923 | 35.385 |
| **Predicted likelihood ≥ 0.5** | 0.000 | 0.000 | 4.615 | 4.549 | 10.454 | 18.742 | 26.000 | 28.812 | 32.862 | 35.374 |
